# Dowsing for salinity tolerance related genes in chickpea through genome wide association and *in silico* PCR analysis

**DOI:** 10.1101/519744

**Authors:** Shaimaa M. Ahmed, A.M. Alsamman, M.H. Mubarak, M.A. Badawy, M.A. Kord, O.A. Momtaz, A. Hamwieh

## Abstract

Soil salinity is a major abiotic stress severely limits agricultural crop production throughout the world, and the stress is increasing particularly in the irrigated agricultural areas. Chickpea (Cicer arietinum L.) is an important grain legume that plays a significant role in the nutrition of the developing world. In this study, we used a chickpea subset collected from the genebank of the International Center for Agricultural Research in the Dry Area (ICARDA). This collection was selected by using the focused identification of germplasm strategy (FIGS). The subset included 138 genotypes which have been screened in the open field (Arish, Sinai, Egypt) and in the greenhouse (Giza, Egypt) by using the hydroponic system at 100 mM NaCl concentration. The experiment was laid out in randomized alpha lattice design in two replications. The molecular characterization was done by using sixteen SSR markers (collected from QTL conferred salinity tolerance in chickpea), 2,500 SNP and 3,031 DArT markers which have been developed and used for association study. The results indicated significant differences between the chickpea genotypes. Based on the average of the two hydroponic and field experiments, seven tolerant genotypes IGs (70782, 70430, 70764, 117703, 6057, 8447 and 70249) have been identified. The data analysis indicated one SSR (TAA170), three DArT (DART2393, DART769 and DART2009) and eleven SNP markers (SNP2021, SNP1268, SNP1451, SNP1487, SNP1667, SNP2095, SNP190, SNP2247 SNP1947, SNP2331 and SNP948) were associated with salinity tolerance. The flanking regions of these markers revealed genes with a known role in the salinity tolerance, which could be candidates for marker-assisted selection in chickpea breeding programs.

## Introduction

About 7.5 billion human share the same land, food and water resources. The global food demands are exponentially expanded while; the water scarcity will affect 1.8 billion people in 2025. One of the most crucial problems that face food security is salinity. According to FAO, over 6.5% of the world’s land is affected, which is translated into 800 million HA of arable lands and expanding dramatically^1^. Filling the gap between the consumption and production require more research in order to enhance unprepared economic plant varieties to face such sudden environmental changes and unlock their ability to tolerance.

Chickpea is one of the candidate cereal crops, which provides food with high nutritional value for an expanding world population. In addition, its global annual production is over 12 million tons. The chickpea production is centered in China (17%), India (12%), Russia and the USA (8%) ^2^. Chickpea is sensitive to salinity which reduces its yield greatly ^3^.

Upon exposure to salt stress, the meristems accumulate salts in the vacuoles of the xylem ^4^, to lower their osmotic potential till reaching high concentrations ^5,6^. Other strategies to tolerate salinity can be by efficient osmotic adjustment, homeostasis, retention in root and mesophyll cells, and ROS detoxification ^7,8^. The sodium (Na^+^) accumulation in the cytoplasm dehydrates the cell by causing ion homeostasis imbalance that inhibits enzyme activity causing cell death and toxicity ^4^. Salinity could affect the activity of the antioxidant enzymes, H2O2 content, chlorophyll fluorescence (Fv/Fm), quantum yield of PSII and the rate of lipid peroxidation in leaf and root tissues ^9^. The morphological effect of salinity on the roots was reported to be agravitropic growth of roots, i.e., growth in diameter rather than length which could reduce plant growth rate by 20%, shoot biomass by 28% and seed yield by 32% ^10,11^.

The complexity of plant gene network makes it hard to procure a single gene with the main responsibility for salinity tolerance in the tolerant cultivars. Although most of the genes clasp with each other through the vast pool of plant biological pathways, triggering one would not mean that there will be no necessity to stimulate the others. A plethora of genes or gene families were reported to have a direct affinity with tolerant plants in response to salinity. Such genes play a role in the physiological or morphological response that the plant system uses as an outlet. Genes such as high-affinity K^+^ transporter (HKT), Na^+^/H^+^ antiporter, Na^+^/Ca^2+^ exchanger are related to ion homeostasis. Serine/threonine protein kinase, peroxidase, calcium-dependent NADPH oxidase gene families are genes related to abiotic stress ^10,12–15^. However, the gene regulatory system could have the higher hand in some cases. The DREB gene family regulates the expression of many stress-inducible genes mostly in an ABA-independent manner in addition to playing a critical role in improving the abiotic stress tolerance of plants by interacting with a DRE/CRT cis-element in the promoter region of various abiotic stress-responsive genes and other TF families ^10,16^.

On the other hand, the massive growth in phenotypic and genotypic assessment technologies has encouraged dissecting genomic loci responsible for crop saline tolerance, in order to improve and enhance cultivated varieties. Researchers’ interest in improving, profiling and mapping salinity tolerance loci has exponentially increased. Classical molecular marker technologies such as simple sequence repeats (SSR) has proved its value in detecting, tagging and identifying salinity tolerance loci in different plant species such as wheat ^17,18^, rice ^19,20^ and chickpea ^21,22^.

Due to their abundance in genomes, evolutionary relationship, suitability for genetic diversity analysis and association with complex phenotypic traits, SNP markers have gained remarkable value in plant molecular genetics ^23^. Genome-wide association study (GWAS) through SNP genotyping has a great impact on identifying genetic regions associated with quantitative and complex traits. GWAS has been used in studying and dissecting genetic construction for salinity tolerance in Rice ^24^, Soybean ^25^, Sesame ^15^, Barley ^26^ and Rapeseed ^27^. The increase in the plant whole genome annotation provides a golden opportunity to weigh trait-associated SNPs according to their potential effects in gene regulation. Such combination between phenotypic and genotypic effects could unravel unseen responsibilities for genomic variations, connect unrelated traits and promotes promising SNPs for high throughput genotyping or genome-editing technologies. Studying trait-associated SNPs was reported in pursuing genes related to 100 seed weight, root/total plant dry weight ratio ^28^, tolerance to herbicide ^29^, tolerance to Ascochyta blight ^30^, tolerance to drought and heat ^31^ in chickpea.

In order to reduce genome complexity, many methods have been developed. The DArT assay provides a remarkable advantage via an inventive selection of genome fraction corresponding predominantly to active genes (http://www.diversityarrays.com/dart-application-dartseq). DArT has been applied for QTL mapping in Rapeseed ^32^, Wheat ^17,33^ and Barley ^34,35^ and for population structure in A. *tauschii* ^36^.

The localization of salinity tolerance related loci in chickpea has been reported in few research articles, such as, ^21^ that reported two key genomic regions on Ca5 and on Ca7, that harboured QTLs for six and five different salinity tolerance associated traits, respectively. Based on the gene ontology annotation, they roughly identified forty-eight putative candidate genes responsive to salinity stress on CaLG05 (31 genes) and on CaLG07 (17 genes). Most of the genes were known to be involved in achieving osmoregulation under stress conditions. In addition, ^10^ used differential expression gene analysis to detect abiotic stress-related genes that significantly were up-regulated in the tolerant genotypes and were down-regulated in the sensitive genotypes under salt stress.

The plant mechanism for both salinity and drought require the activation of genes responsible for the plant cellular adaptation, the cell wall maintenance, the protection against water loss and the cellular ionic and osmotic homeostasis ^37^. This relationship between salinity and drought was employed in different plant molecular genetics researches, thus drought tolerance responsible loci could be used to expand the research in salinity and vice versa ^38–41^. In a breeding Chickpea population resulted from a cross between ICC4958 and Annigeri, two loci on LG4, were found to be associated to drought tolerance ^42^.

In order to, improve the efficiency with which specific adaptive traits are identified from genetic resource collections the Focused Identification of Germplasm Strategy (FIGS) was designed based on the premise that the environmental selection pressures from which these germplams was originally sampled will be reflected on them ^43^. FIGS uses both trait and environmental data to define a set of accessions with a high probability of containing the desired traits based on a quantification of the trait-environment relationship ^44–46^. FIGS has been successfully used to screen new genes related to abiotic and biotic stresses in different plant species ^43,47–49^.

In the present study, authors used FIGS protocol to construct a chickpea diversity panel that contains landraces and wild chickpea accessions with a potential salt stress tolerance. In addition to comparing the performance of one dominant (DArT) and two co-dominant (SSR and SNP) genetic markers in studying the relatedness coefficients between these genotypes, population patterns and gene diversity. The molecular markers associated with chickpea salinity tolerance were studied according to their genetic effect and their gene closeness through SNP effect and *In silico* PCR analysis. Previously published salinity and drought tolerance associated SSR markers were studied through *In silico* PCR analysis in order to identify closed genes. All the potential genes were studied according to their gene pathways and the relatedness to chickpea salinity tolerance.

### Plant Material

The studied germplasm panel is composed of 203 different genotypes that were collected from 28 provinces in 13 countries across the globe. The seeds were provided by the gene bank of ICARDA by using FIGS tool (TABLE S1). Most of the chickpea panel is from Pakistan and India provinces which are thought to have saline environments. 138 genotypes were used in both the hydroponic and field environments.

### Salinity tolerance of chickpea genotypes

The experiments were performed in the field (Arish, Sinai, Egypt) and in the greenhouse by using the hydroponic system (Agricultural Research Centre, Giza, Egypt), in November 2014. The individuals were replicated two times and arranged in an alpha lattice randomized block design. A dripping water irrigation system was installed to water the field every two weeks. In the greenhouse, one seed of individual accession was sown in a tray (10 cm - 30 cm) containing a mixture of peat moss (40%) and perlite (60%). After two weeks of germination, the seedlings were transferred to hydroponic tank (150 cm _ 230 cm) with half strength of a nutrient solution.

To determine the salt concentration in the soil of the field, a soil sample was air dried, softened and sieved prior to preparing soil pest then soil solution was extracted to determine pH, and the concentration of cations and anions (Page, 1982). The salt concentration of the hydroponic tank was adjusted at 100 mM and the pH was adjusted at 8, both of them were checked daily. Salinity stress tolerance trait was evaluated as the necrosis score. The scoring was conducted at a relatively early stage of salt treatment when the necrotic symptoms appeared on the stem and the leaves, after one week. The phenotypic scaling was set from 1 to 5, where the tolerant plant was given score number 1, and the sensitive plant was given score number 5. And when the plant is partly tolerant, it was given number 2, 3 or 4 according to the intensity of the necrosis. The phenotypic reading of the greenhouse was taken every three weeks during the experiment time. For the field experiments, the phenotypic readings were taken every two months. One way ANOVA was performed to test the significance of both the field and the greenhouse phenotypic data. Consequently, the average data was used for further analysis. Figure 4 shows the frequency distribution of salinity tolerance among plant genotypes.

**Figure 1:**
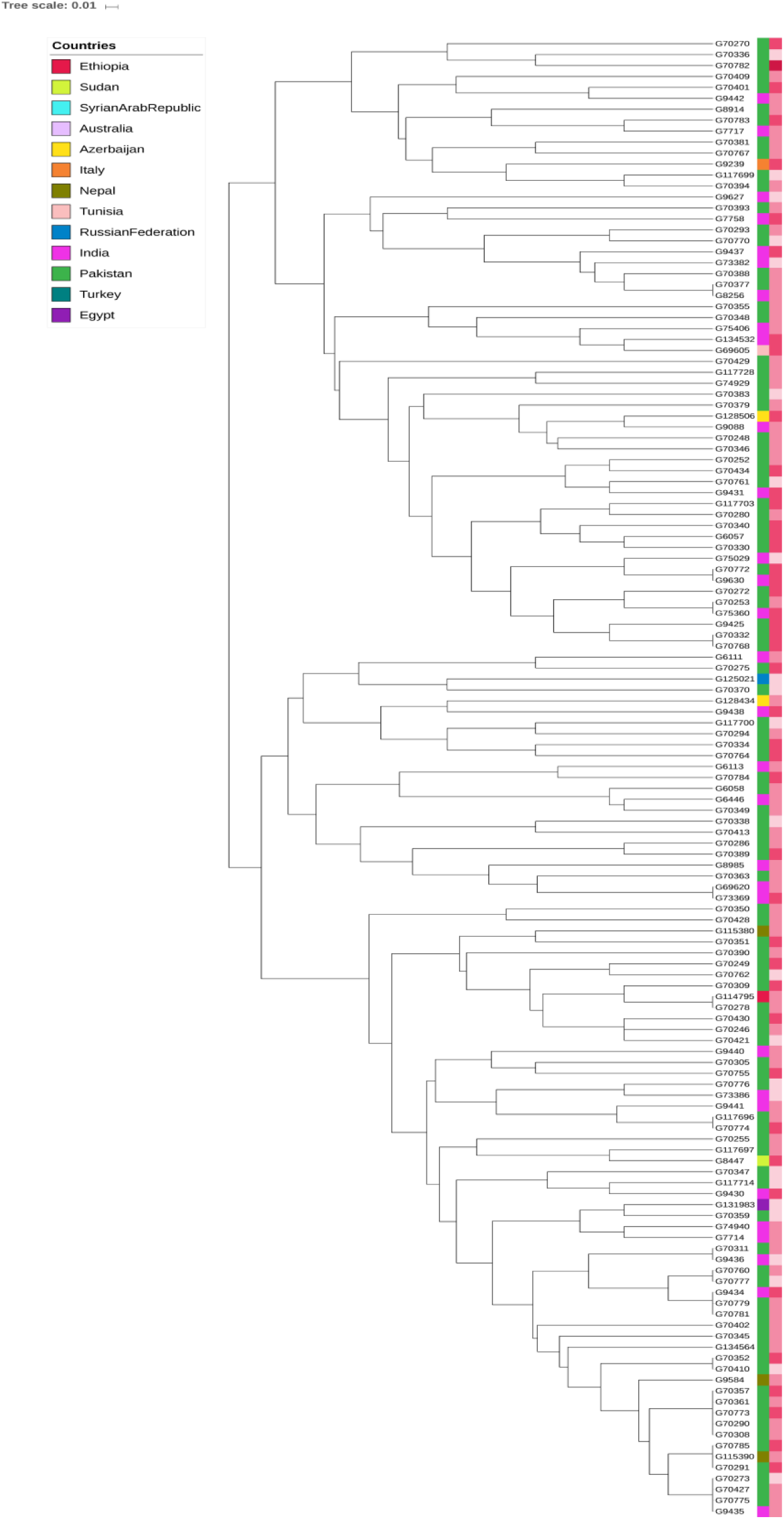
The phyolegenetic tree of chickpea genotypes based on SSR markers.

**Figure 2:**
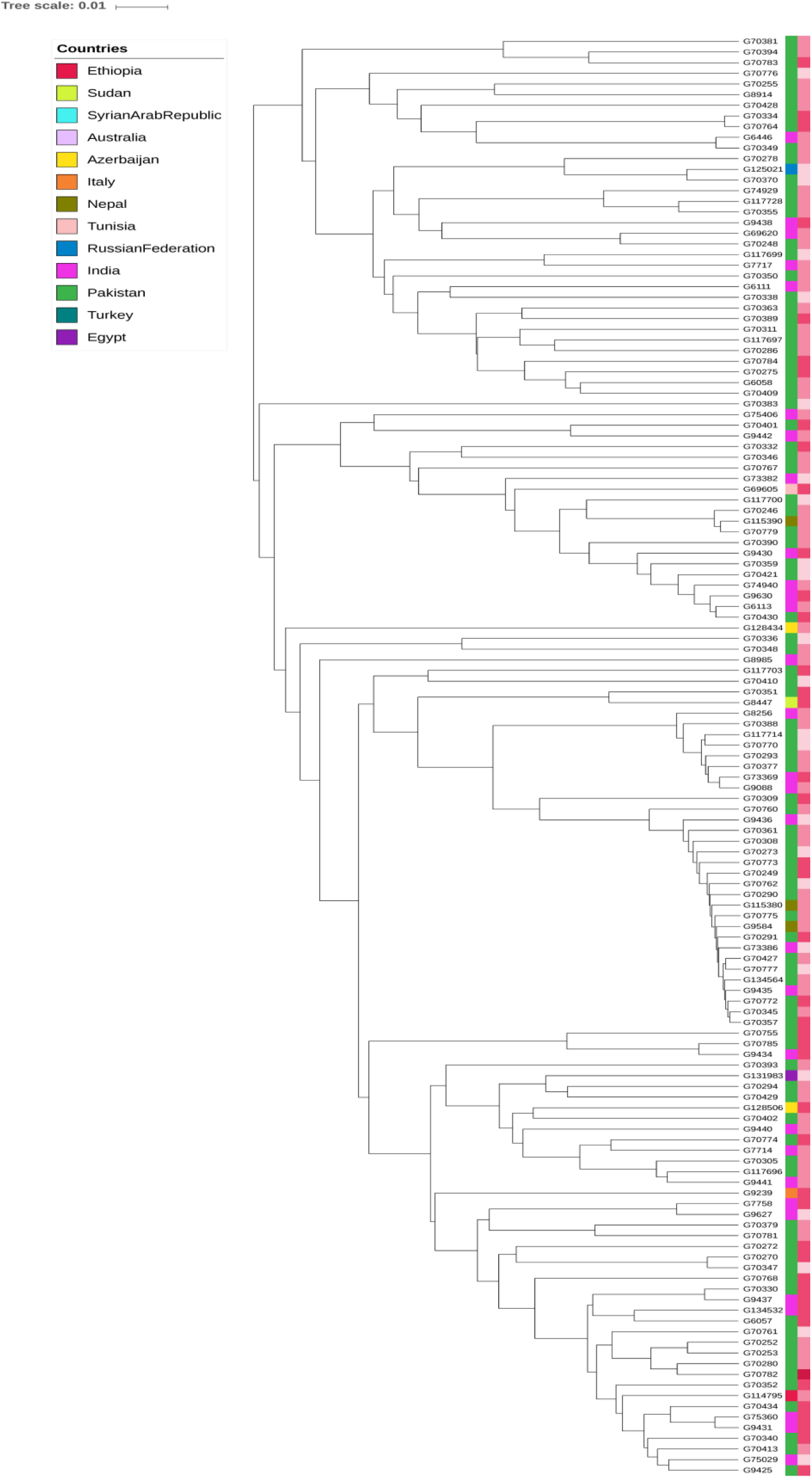
The phyolegenetic tree of chickpea genotypes based on DArT markers.

**Figure 3:**
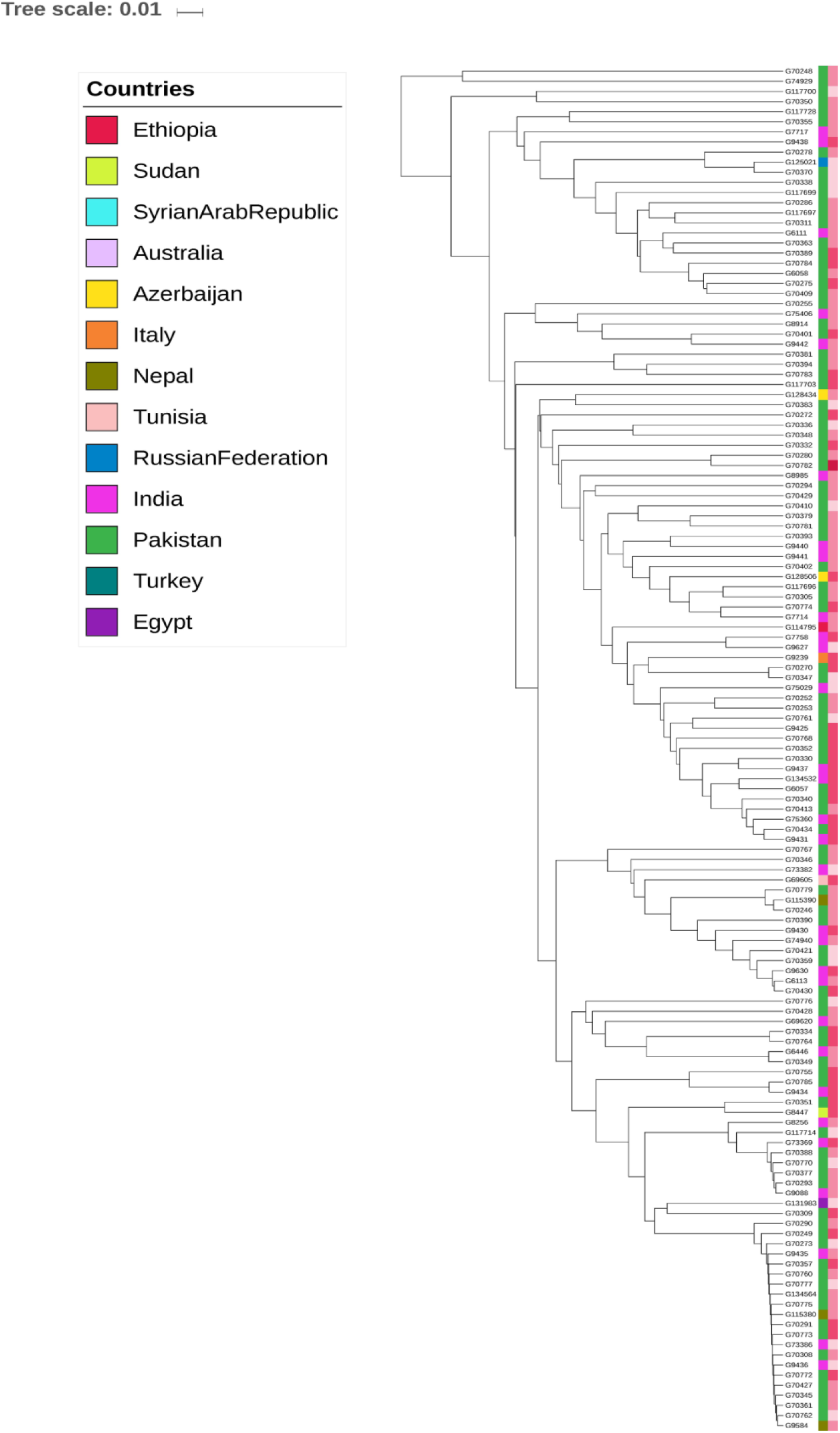
The phyolegenetic tree of chickpea genotypes based on SNP markers.

**Figure 4:**
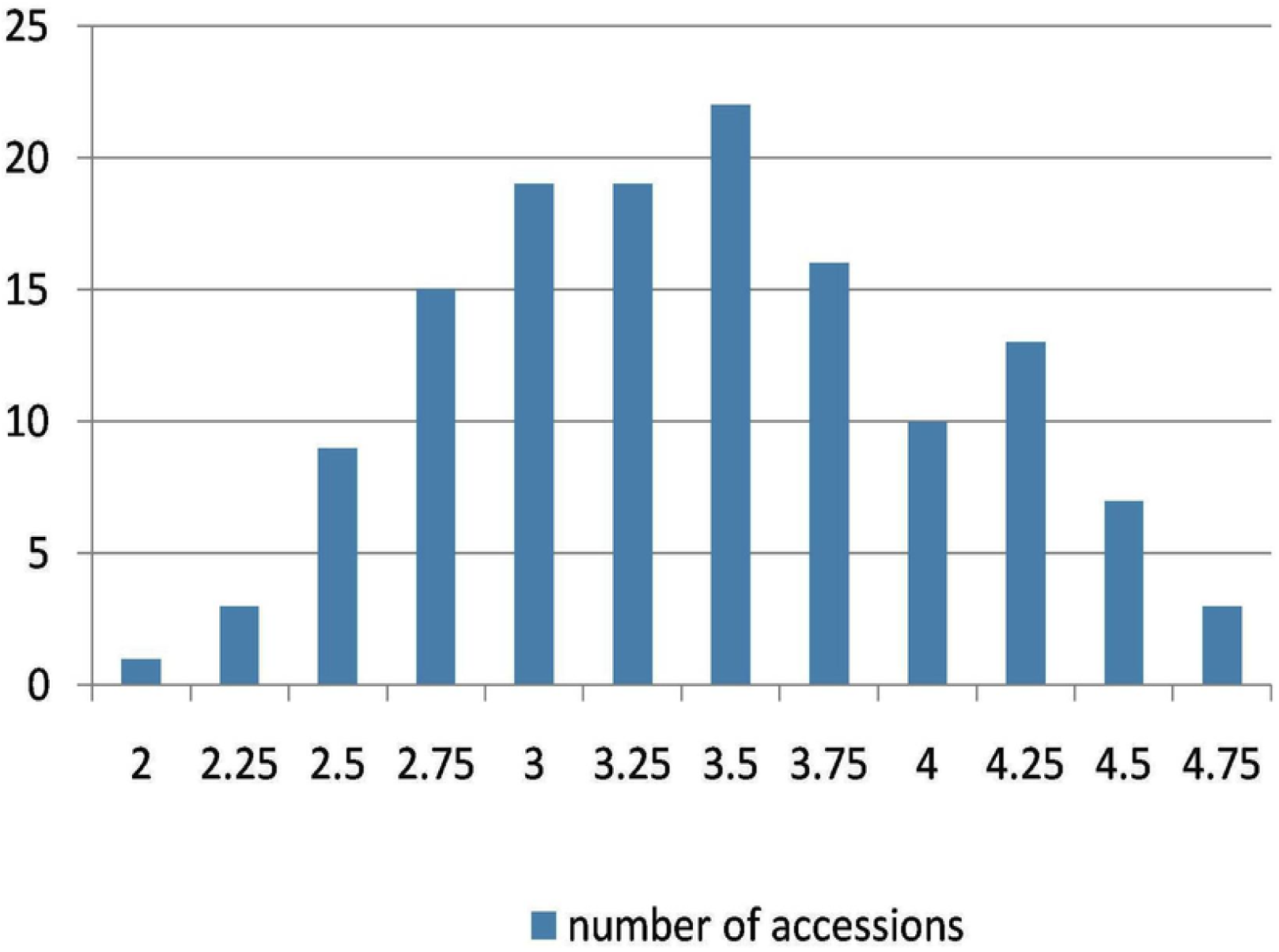
The phenotypic distribution for salinity tolerance among chickpea genotypes.

### DNA Extraction

About 0.1 gm. of fresh tissue was ground by liquid nitrogen using mortar and pestle. Then 1 ml. CTAB was added to the ground samples and mixed well to be incubated for 1 hour at 65 °C. 1 ml. chloroform: Isoamyl (24:1) was added to the samples and the mixture was shaken for 20 minutes. Then the samples were centrifuged for 15 minutes, and the supernatant was transferred to a new tube. 1 ml of absolute cold isopropanol was used to precipitate the DNA. Then the samples were centrifuged. The pellet was washed twice with 70% ethanol. The samples were air-dried and eluted in 200 µl 1X TE ^50^.

### Molecular Markers

Sixteen SSR primers have been used in this study. TABLE 1 represents the review of literature of the polymorphic SSR markers. SSR PCR reactions were performed in 15 µl reaction volume consisting of 5 ng DNA template, 10 picomol of forward primer, 10 picomol of reverse primer, 0.1 U of Taq DNA polymerase, 25 mM of MgCl2, 2 mM dNTPs and 10X PCR buffer in 96-well microtitre plates using thermal cycler. PCR program was used to amplify DNA fragments as follow: initial denaturation was 5 min at 95°C. This was followed by 35 cycles of denaturation for 15 sec at 95°C, annealing for 15 sec at 55°C and extension for 30 sec at 72°C. Subsequently, 7 min final extension at 72°C. SSR markers were checked for amplification on 9 % acrylamide gel.

**Table 1:**
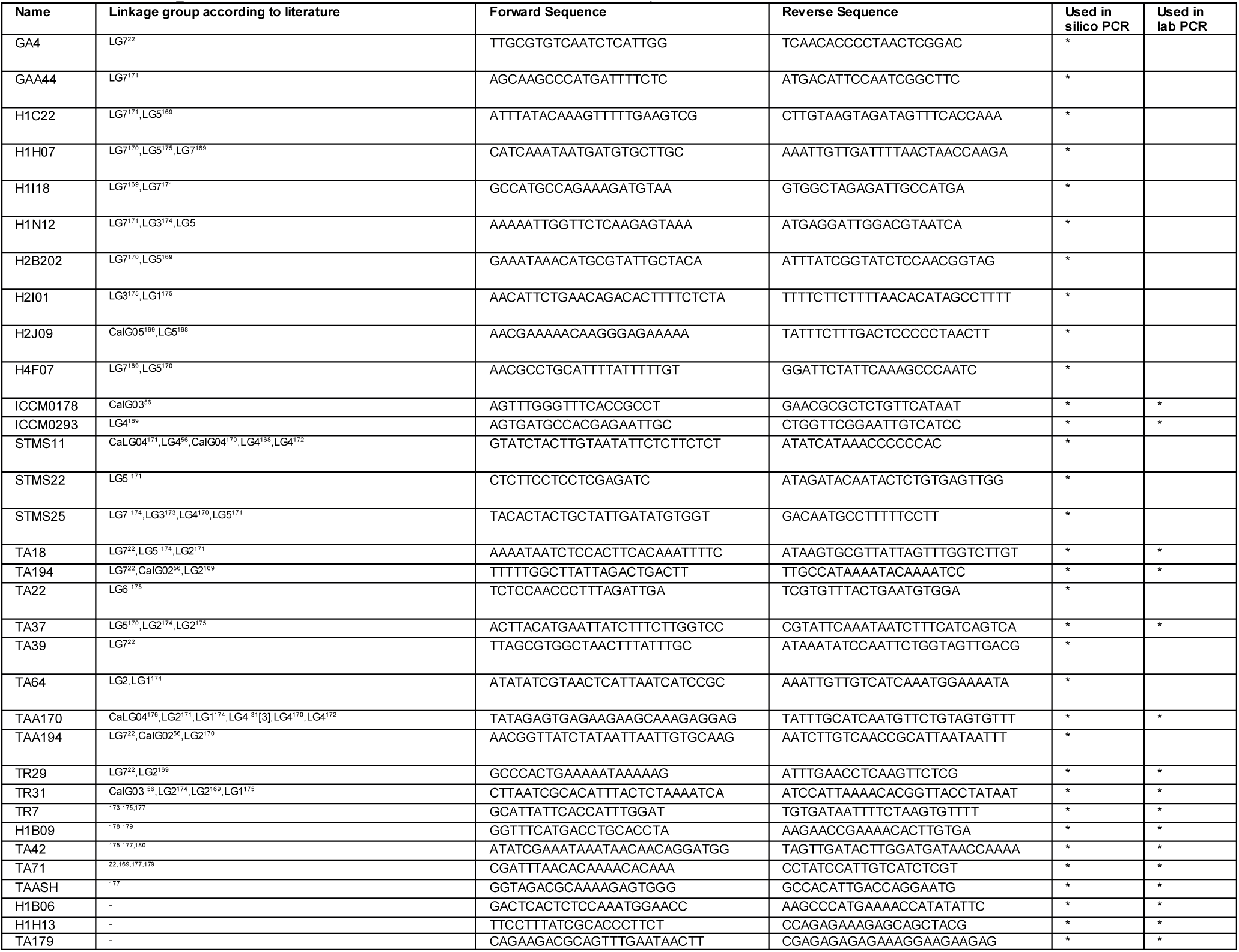
SSR primers’ information used in this study.

The Diversity Arrays Technology (DArT®) markers panel was used to genotype chickpea population with high-density. Out of 203 different genotypes that were collected, 186 genotypes were sent for DArT, using 50 µl of a 100 ng µl/1 DNA of each sample. The DNA was sent to Triticarte Pty. Ltd. Australia (http://www.triticarte.com.au) for DArT markers genotyping (Chickpea DArTseq panel version 1.0) and SNP genotyping as a provider for commercial service. A 3031 DArT and 2499 SNP polymorphic marker loci with quality parameter and call rate both greater than 80 % and minor allele frequency (MAF) > 5 % were selected for genome-wide association analysis.

BLAST tool ^51^ was used to assign SNP and DArT markers to chickpea chromosomes. DArT markers were assigned to all chickpea chromosomes, where Ca4 has the highest number of DArT markers (755), Ca8 has the lowest number (127) and 69 markers has unknown position (Ca9). For SNP, Ca4 has the highest number of markers (632), while Ca8 has the lowest number (127) and 13 makers with unknown location (Ca9). The marker density in DArT assay ranged from 17 (Ca6) to 54 markers/Mpb (Ca3), while in SNP assay it ranged from 19 (Ca8) to 48 (Ca7) marker/Mpb (Ca4) (TABLE 2).

**Table 2:** Genomic distribution of SNPs and DArTs physically mapped on eight chickpea chromosomes.

| Chromosome | DArT marker count | DArT Density | SNP Density | SNP Marker count |
| --- | --- | --- | --- | --- |
| Ca1 | 374 | 19 | 26 | 333 |
| Ca2 | 243 | 23 | 27 | 233 |
| Ca3 | 277 | 25 | 31 | 236 |
| Ca4 | 755 | 27 | 35 | 632 |
| Ca5 | 290 | 33 | 35 | 244 |
| Ca6 | 483 | 17 | 41 | 376 |
| Ca7 | 413 | 49 | 48 | 305 |
| Ca8 | 127 | 54 | 19 | 127 |

### Molecular marker analysis

Genetic distances (GD) and Nei’s gene diversity ^52^, were calculated for each locus separately for every marker type. Based on the GD matrices, phenograms of the 186 samples were constructed with the Unweighted Pair Group Method with Arithmetic means (UPGMA) by Power marker software ^53^. The association between SSR markers and salinity stress tolerance was performed through F-test for each marker to test its linkage to the salinity stress tolerance through PowerMarker software ^53^. Spoon *in silico* PCR program (http://www.ageri.sci.eg/index.php/facilities-services/ageri-softwares/spoon) was used to identify the location of SSR marker loci previously published and used through study, these SSR markers are known to be linked to salinity or drought tolerance in chickpea. While, The association of the SNP and the DArT markers with salinity stress tolerance was performed by using GAPIT (R Package) ^54^. SNP and DArT markers which have been used for further analysis, have a significance score higher than higher than 0.0001 in both open field and greenhouse. The effect of SNPs linked to salinity tolerance has been studied through SnpEFF ^55^. The chickpea genome ^56^ has been used through all marker-gene analysis. Genes near/adjoin salinity associated molecular markers have been used for gene enrichment analysis, the gene pathways for these genes have been determined by KEGG pathways database ^57^ using BlastKOALA tool ^58^. Circos configuration ^59^ was used to illustrate the location of the significant molecular markers and candidate genes, from which the distance between the QTL and the gene can be assumed. The iTOL online web tool ^60^ was used to draw phylogenetic trees, while ClustVis ^61^ was used to construct population structure kinship illustration.

## Discussion

Chickpea is widely grown in West and Central Asia and Australia, where saline soils are abundant. There is narrow genetic variation among different genotypes, which is an obstacle for breeding for salinity tolerance. In chickpea, despite the conductance of several mapping studies, only two studies have reported the presence of QTLs for salinity tolerance. There is no report on putative candidate genes that would confer salinity tolerance in chickpea ^21^.

### Soil analysis of field experiment in Arish

The chickpea evaluations for salinity tolerance were made across the years 2014 and 2015 through field and greenhouse experiments. The field trial was made in the open field (Arish, Sinai, Egypt). The salt concentration in the field was 344 ppm in the first 30 cm depth, 904 ppm in the depth from 30 to 60 cm, and 848 ppm in the depth of more than 60 cm. The analysis of the water of irrigation revealed that the average of the salt concentration was 897 mM, and the pH was 7.2.

### Phenotypic analysis for salinity tolerance

The plant which takes reading till 3.25 was considered to be tolerant. The plant which takes 5 as necrosis scale was considered to be highly sensitive (died by stress). Forty-seven accessions were observed to be tolerant and 3 accessions were highly sensitive to salinity stress. The salinity tolerance was normally distributed among chickpea genotypes (FIGURE 4). A 138 genotypes were successfully screened in the field and the greenhouse (Arish, Egypt and hydroponic). No significant difference has been observed between the two methods. Significant differences have been identified between the genotypes where CV% and LSD was 7.8 and 1.27, respectively.

The phenotypic evaluation showed significant variations for salinity stress tolerance under saline conditions between various genotypes indicating a broad phenotypic variance within the global chickpea population. Based on the average from the two hydroponic and field experiments, seven tolerant genotypes IGs (70782, 70430, 70764, 117703, 6057, 8447 and 70249) have been identified (TABLE 3). The present data indicated the presence of genetic components influencing salinity tolerance traits. Using ANOVA, we found significant constancy of the salinity stress tolerance trait suggesting the genetic control of this trait as seen in (TABLE 4). It was observed from the tables that among the seven most tolerant genotypes, six of them are from Pakistan, and one from Azrabejan.

**Table 3:** The most salinity tolerant genotypes obtained by analysis of variance (ANOVA) from the phenotypic readings of the green house and the field.

| Genotype | Field | Hydroponic | Origin | Province |
| --- | --- | --- | --- | --- |
| IG70782 | 2 | 2 | Pakistan | Punjab |
| IG70430 | 2.5 | 2 | Pakistan | Punjab |
| IG70764 | 2.5 | 2 | Pakistan | Sindh |
| IG117703 | 2.5 | 2 | Pakistan | Punjab |
| IG6057 | 2.5 | 2.5 | Pakistan | NWF |
| IG8447 | 2.5 | 2.5 | Azrabejan | Lankaran |
| IG70249 | 2.5 | 2.5 | Pakistan | Sindh |

**Table 4:** The ANOVA output for differences between the green house and field phenotyping.

| Source of variation | d.f. | SS | MS | F |
| --- | --- | --- | --- | --- |
| Genotype | 137 | 210.88 | 1.54 | 1.86*** |
| Location | 1 | 1.04 | 1.04 | 1.26 |
| Genotype X Location | 137 | 150.46 | 1.1 | 1.33 |
| Error | 275 | 227.41 | 0.83 |  |
| Total | 551 | 609.38 |  |  |

### Simple sequence repeat analysis

Polymorphic SSRs are excellent molecular markers, because of their multi-allelism, genome abundance, co-dominance and high polymorphic rate. Additionally, compared to high throughput molecular markers such as SNPs, SSRs have their own advantages respected to population genetics ^62^. The association between SSR markers and abiotic stresses tolerance such as salinity has been recognized and discussed in different plant species ^39,63^. One of the main importance of the SSR markers, the inheritance of its alleles from the parents to the progeny which made them an efficient tool in marker-assisted selection breeding programs.

Sixteen SSR markers were applied in this study, these markers have shown linkage to salinity tolerance from previous literature (TABLE T1). The polymorphism information content (PIC) detect the ability of the marker to find the genetic variation among the used diversity set. In addition, the PIC is a measure of the marker informativeness and it ranges from 0 to 1. The markers with a PIC higher than 0.5 are highly informative, while, a PIC value between 0.5 and 0.25 implies a locus of moderate informativeness ^64^. The PIC value in the present study ranged from 0.3 (H1H13) to 0.9 (TAA170).

The marker-trait association analysis for SSR markers was calculated separately using greenhouse and field data. Additionally, the analysis of variance (ANOVA) was calculated for greenhouse and field results. The significance score for the association between SSR markers abundance and greenhouse data, ranged from 0.006 (TAA170) to 1 (TA179), while with field data, it ranged from (0.06) (TAA170) to 0.9 (TR31), with ANOVA score ranged from 0.02 (TAA170) to 0.9 (TA179). The significant association between TAA170 and salinity stress tolerance indicates a potential role in saline tolerance control. Table 5 represents the sixteen polymorphic SSR markers used in this study and their association with the salinity stress tolerance in the hydroponic experiment in the greenhouse and Arish’s field environments.

**Table 5:** The 16 polymorphic SSR markers used and their association to the salinity stress tolerance in the hydroponic experiment in the green house and Arish field environments.

| Marker | PIC | hydroponic | Arish field | ANOVA |
| --- | --- | --- | --- | --- |
| ICCM0178 | 0.5 | 1 | 0.7 | 0.4 |
| TAA170 | 0.9 | 0.006 | 0.06 | 0.02 |
| H1H13 | 0.3 | 0.6 | 0.8 | 0.8 |
| H1B06 | 0.5 | 0.3 | 0.4 | 0.9 |
| TA194 | 0.7 | 0.3 | 0.4 | 0.07 |
| TA37 | 0.5 | 0.2 | 0.6 | 0.3 |
| TA18 | 0.8 | 0.8 | 0.1 | 0.6 |
| ICCM0293 | 0.7 | 0.9 | 0.4 | 0.9 |
| TA42 | 0.8 | 0.2 | 0.4 | 0.6 |
| TR31 | 0.6 | 0.5 | 0.9 | 0.4 |
| TA179 | 0.7 | 1 | 0.4 | 0.9 |
| TR29 | 0.8 | 0.52 | 0.2 | 0.2 |
| TA71 | 0.7 | 0.2 | 0.4 | 0.1 |
| TAASH | 0.6 | 0.4 | 0.7 | 0.5 |
| H1B09 | 0.7 | 0.1 | 0.4 | 0.6 |
| TR7 | 0.6 | 0.6 | 0.6 | 0.5 |

SSR phylogenetic analysis clustered chickpea genotypes into two separated clusters. On the origin level, SSRs were almost successful to combine Pakistani and Indian genotypes. Most of the tolerant chickpea genotypes were clustered in the first clusters, while moderately tolerant genotypes were clustered in the second major cluster (FIGURE1).

*In silico* PCR is a computational procedure that estimates PCR results theoretically using a given set of primers to amplify DNA sequences from a sequenced genome or transcriptome ^65,66^. This procedure could offer the ability to explore and detect QTLs nearest genes, in order to study their potential relationship with traits of interest. Several QTLs which have been acquired through SSR markers, showed a significant relationship with abiotic stresses such as salinity and drought in chickpea, where the association between these loci and their corresponding traits was presented through field trials and lab experiments. In order to study and explore chickpea genes nearby our and previously published chickpea salinity tolerance QTLs, the *in silico* PCR analysis was conducted on our lab-tested SSR markers in addition to previously published chickpea SSR markers to study and explore the chickpea nearby genes (TABLE 1).

*In silico* PCR analysis revealed that 15 SSR markers were closely located near 19 chickpea genes with a distance ranged from 6 bp to 7 kbp (TABLE 6). According to UniProtKB (www.uniprot.org), which is a central hub for the collection of functional information on proteins, the gene ontology analysis showed that, at the molecular function level some of these genes belong to phosphorelay sensor kinase activity (1), DNA binding transcription factor activity (1), catalytic activity (12) and binding (8). While, at the cellular component level, some shows a relationship to Golgi membrane (1), mitochondrial matrix (1), integral component of membrane (5), protein-containing complex (2), organelle (1), organelle part (2) and cell part (5). Additionally, on the biological process level some genes belong to metabolic process (4), cellular process (4) and cell wall organization (1).

**Table 6:** The SNPs, DArTs and SSR markers close genes and gene effect.

| Chromosome | Start | End | Gene name | SNP Name | Start | End | Effect | Marker Type |
| --- | --- | --- | --- | --- | --- | --- | --- | --- |
| Ca1 | 22102973 | 22103524 | NIPBL | SNP1268 | 3934547 | - | MODIFIER | SNP |
| Ca1 | 6377188 | 6381838 | UN | SNP1451 | 22106206 | - | MODIFIER | SNP |
| Ca1 | 6386498 | 6388905 | MAP-NDL | SNP1487 | 6383629 | - | MODIFIER | SNP |
| Ca2 | 3920563 | 3935917 | P67 | SNP1487 | 6383629 | - | MODIFIER | SNP |
| Ca3 | 39213565 | 39216469 | PUB9 | SNP1667 | 39215603 | - | MODIFIER | SNP |
| Ca3 | 14517059 | 14520739 | TOMM6 | SNP190 | 14518700 | - | MODIFIER | SNP |
| Ca3 | 23326348 | 23329856 | TT12 | SNP1947 | 23327600 | - | MODIFIER | SNP |
| Ca3 | 23331054 | 23334408 | E3_ubiquitin_ligase | SNP1947 | 23327600 | - | MODIFIER | SNP |
| Ca4 | 45624806 | 45631675 | APEX2 | SNP2021 | 14771678 | - | MODIFIER | SNP |
| Ca4 | 16528933 | 16529810 | MED4 | SNP2021 | 14771678 | - | MODIFIER | SNP |
| Ca4 | 16529838 | 16533251 | random_slug_p5 | SNP2095 | 13746772 | - | MODIFIER | SNP |
| Ca4 | 16535273 | 16538805 | VPS39 | SNP2095 | 13746772 | - | MODIFIER | SNP |
| Ca4 | 13717510 | 13717602 | kif11 | SNP2331 | 45628242 | - | MODIFIER | SNP |
| Ca5 | 19845561 | 19857555 | UN | SNP948 | 16530812 | - | MODIFIER | SNP |
| Ca7 | 14767114 | 14772777 | HBP-1b(c38) | SNP948 | 16530812 | - | MODIFIER | SNP |
| Ca7 | 14774632 | 14777337 | CTL | SNP948 | 16530812 | - | MODIFIER | SNP |
| Ca7 | 13738211 | 13742001 | HIS3 | DART2009 | 13717660 | - | UNKNOWN | DART |
| Ca7 | 13749560 | 13759327 | HIBCH | DART769 | 19859119 | - | UNKNOWN | DART |
| Ca1 | 8579323 | 8579615 | HYPK | GAA44 | 8579440 | 8579624 | UNKNOWN | SSR |
| Ca1 | 8579323 | 8579615 | PBL | GAA44 | 8582087 | 8589091 | UNKNOWN | SSR |
| Ca1 | 30480316 | 30480448 | UC | TA22 | 30481063 | 30484174 | UNKNOWN | SSR |
| Ca1 | 30480316 | 30480448 | UC | TA22 | 30481197 | 30481421 | UNKNOWN | SSR |
| Ca1 | 280162 | 280372 | EIF3C | TAA170 | 14148718 | 14153238 | UNKNOWN | SSR |
| Ca1 | 280162 | 280372 | ASK10 | TAA170 | 14155412 | 14156240 | UNKNOWN | SSR |
| Ca1 | 280162 | 280372 | CHSP70 | TAA170 | 267731 | 271187 | UNKNOWN | SSR |
| Ca2 | 7517182 | 7518107 | USP17L2 | H1N12 | 46943791 | 46944124 | UNKNOWN | SSR |
| Ca2 | 7517182 | 7518107 | UC | H1N12 | 7501062 | 7507760 | UNKNOWN | SSR |
| Ca2 | 32200882 | 32201085 | Cucumisin-like | TA194 | 16817161 | 16817183 | UNKNOWN | SSR |
| Ca2 | 17196291 | 17196573 | PERK9 | TA37 | 17186142 | 17192315 | UNKNOWN | SSR |
| Ca3 | 12965840 | 12965951 | TFIID | H2J09 | 27319090 | 27322337 | UNKNOWN | SSR |
| Ca3 | 16591364 | 16591641 | RPL32 | ICCM0178 | 16592795 | 16594742 | UNKNOWN | SSR |
| Ca3 | 16591364 | 16591641 | OXI1 | ICCM0178 | 16585565 | 16588533 | UNKNOWN | SSR |
| Ca3 | 16819768 | 16819858 | bHLH62 | TA194 | 45762276 | 45767196 | UNKNOWN | SSR |
| Ca3 | 29597808 | 29598023 | DREB2F | TA64 | 29600041 | 29601190 | UNKNOWN | SSR |
| Ca3 | 29597808 | 29598023 | SMUBP-2 | TA64 | 29601398 | 29608471 | UNKNOWN | SSR |
| Ca3 | 29597808 | 29598023 | UC | TA64 | 29575704 | 29593101 | UNKNOWN | SSR |
| Ca3 | 29597808 | 29598023 | UC | TA64 | 29592767 | 29592949 | UNKNOWN | SSR |
| Ca3 | 3477462 | 3559535 | RPP1C | TAA194 | 3557121 | 3557498 | UNKNOWN | SSR |
| Ca3 | 3477462 | 3559535 | UC | TAA194 | 3481336 | 3481777 | UNKNOWN | SSR |
| Ca4 | 9102502 | 9102731 | DLAT | CaSTMS11 | 9102461 | 9102521 | UNKNOWN | SSR |
| Ca4 | 9102502 | 9102731 | PK | CaSTMS11 | 9104818 | 9110326 | UNKNOWN | SSR |
| Ca4 | 8913305 | 8914136 | CKI1 | ICCM0293 | 8912696 | 8913273 | UNKNOWN | SSR |
| Ca4 | 8913305 | 8914136 | UC | ICCM0293 | 8917824 | 8920834 | UNKNOWN | SSR |
| Ca4 | 14146314 | 14146529 | CRK25 | TAA170 | 273319 | 279318 | UNKNOWN | SSR |
| Ca4 | 14146314 | 14146529 | PR | TAA170 | 14138672 | 14139743 | UNKNOWN | SSR |
| Ca4 | 14146314 | 14146529 | SYP24 | TAA170 | 289201 | 293996 | UNKNOWN | SSR |
| Ca5 | 35796257 | 35796416 | trc | CaSTMS22 | 35796138 | 35796280 | UNKNOWN | SSR |
| Ca5 | 35796257 | 35796416 | SNRPA | CaSTMS22 | 35800401 | 35803070 | UNKNOWN | SSR |
| Ca5 | 38745589 | 38745797 | ABHD17B | GA4 | 38745778 | 38746079 | UNKNOWN | SSR |
| Ca5 | 38745589 | 38745797 | tipD | GA4 | 38736360 | 38740931 | UNKNOWN | SSR |
| Ca5 | 27314665 | 27314754 | eff | H2J09 | 27310193 | 27310262 | UNKNOWN | SSR |
| Ca5 | 27314665 | 27314754 | UC | H2J09 | 12972787 | 12980678 | UNKNOWN | SSR |
| Ca5 | 9535702 | 9535872 | MOCS2 | H4F07 | 9535822 | 9537098 | UNKNOWN | SSR |
| Ca5 | 9535702 | 9535872 | MAIL3 | H4F07 | 9540227 | 9540326 | UNKNOWN | SSR |
| Ca5 | 23840621 | 23841280 | UC | TA18 | 23787120 | 23831853 | UNKNOWN | SSR |
| Ca5 | 23840621 | 23841280 | UC | TA18 | 23831565 | 23831853 | UNKNOWN | SSR |
| Ca5 | 10350148 | 10350439 | 2014-03-03 | TA39 | 10344830 | 10346113 | UNKNOWN | SSR |
| Ca5 | 10350148 | 10350439 | UC | TA39 | 10339812 | 10340291 | UNKNOWN | SSR |
| Ca6 | 46936179 | 46937651 | AMPD | H1N12 | 7519343 | 7521986 | UNKNOWN | SSR |
| Ca6 | 41372711 | 41373830 | 7OMT | TA18 | 41381250 | 41381294 | UNKNOWN | SSR |
| Ca7 | 37936821 | 37936997 | BON1 | CaSTMS25 | 37936934 | 37937074 | UNKNOWN | SSR |
| Ca7 | 45756763 | 45757560 | rl5 | TA194 | 32201780 | 32202313 | UNKNOWN | SSR |

SSR markers illustrated a close distance to some important chickpea genes; most of these genes have a direct or indirect relationship to salinity and drought tolerance. These genes included DLAT, ASK8, GAUT1, MOCS2, OXI1, CKI1, PERK9, DREB2F, STN7, TMED2 and CRK25 (TABLE6 and FIGURE5).

**Figure 5:**
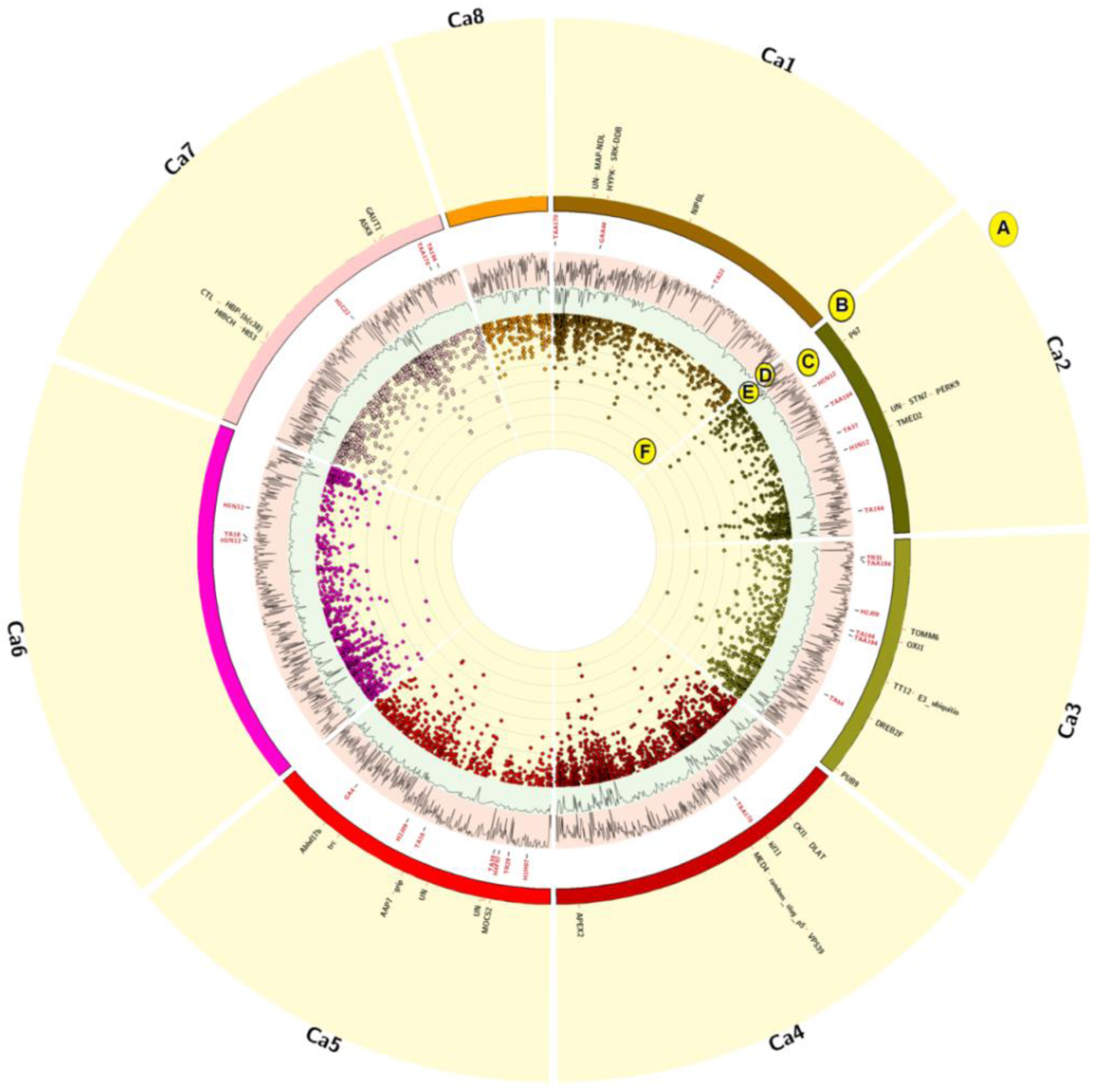
Circos configuration depicting the distribution of genes (B) associated with SSR (C), SNP and DArT markers in addition to their P-value (−log10) scores, SNP markers density (E) and whole genome genes density (D).

DLAT (Dihydrolipoyllysine-residue acetyltransferase) is a component of pyruvate dehydrogenase complex (PDC) in mitochondria and a member of pyruvate metabolic pathway. The PDC catalyzes the overall conversion of pyruvate to acetyl-CoA and CO2 and thereby links the glycolytic pathway to the tricarboxylic cycle ^67^. Taylor *et al*., ^68^reported that environmental stress causes oxidative damage to the mitochondria leading to inhibition of glycine decarboxylase. The participation of pyruvate dehydrogenase in plant response to abiotic stresses was reported in drought, chilling and salinity stresses ^69^.

The salinity stress could affect the plant growth and development ^70^. ASK8 or ASKθ which belongs to the SHAGGY/GSK-3 family plays an important role in serine/threonine kinase activity, which is involved in cell differentiation. The SHAGGY/GSK-3 family was strongly suggested to be involved in stress responses and ASKθ may have a role in the regulation of transcription factors in Arabidopsis ^71^. Recent studies have revealed that plant GSK3 proteins are actively implicated in hormonal signaling networks during development as well as in biotic and abiotic stress responses, especially in osmotic stress responses ^72,73^. Additionally, the over expression of GSK1 influences NaCl stress responses in the absence of NaCl stress and develops enhanced tolerance NaCl tolerance in Arabidopsis ^74^.

Galacturonosyltransferase (GAUT) is a type of α-1,4-galacturonosyltransferase that can transfer galacturonic acid from uridine 5′-diphospho-galacturonic acid into the pectic polysaccharide homogalacturonan ^75^. Reported mutations in the genes belonging to GAUT family resulted in discernible changes in cell wall monosaccharide composition ^76^. GAUT1 and GAUT7 are the main components of cell wall pectin biosynthetic homogalacturonan galacturonosyltransferase complex in plants, where GAUT1 is involved in homogalacturonan (HG) synthesis ^77^.

Molybdenum cofactor (MoCo) biosynthesis is a highly conserved biochemical pathway resulting in the biochemical activation of molybdenum after binding the dithiolene moiety of a small organic compound called molybdopterin ^78^. MOCS2 is the catalytic subunit of the molybdopterin synthase complex and acts as a sulfur carrier required for molybdopterin biosynthesis ^79–81^. It was found that Mo deficiency affects plant nitrogen and sulphur metabolisms, in a manner similar to the nitrate assimilation activity which is inhibited by the shortage of Mo-co, but still different from nitrate or sulphate limitation ^80^.

In response to stimuli and during development of active oxygen species (AOS), genes are generated to function as signalling molecules in order to generate specific downstream responses in eukaryotes ^82^. In A. thaliana, OXI1 kinase is necessary for oxidative burst-mediated signalling in stress response, where OXI1 encodes serine/threonine kinase which is induced in response to a wide range of H2O2-generating stimuli ^82,83^. Additionally, OXI1 controls singlet oxygen-induced cell death in A.thaliana under high-light stress ^84^ and is up-regulated in fungus-infected A.thaliana roots ^85^.

In order to regulate female gametophyte development and vegetative growth in Arabidopsis, the histidine kinase CKI1 acts upstream of histidine phosphotransfer proteins ^86^. On the other hand, the putative Arabidopsis sensor histidine kinases were originally classified as a candidate cytokinin receptor, based on the fact that overexpression of CKI1 in Arabidopsis hypocotyl segments resulted in callus proliferation and shoot differentiation in the absence of exogenously supplied cytokinin and auxin ^87^. There is structure and binding specificity of the receiver domain of sensor histidine kinase CKI1 in A.thaliana where, there are three cytokinin receptors, HK2, HK3 and HK4/CRE1, one putative osmosensing HK, HK1, and two cytokinin independent HKs, CKI1 and CKI2/AHK5 ^88^. In Arabidopsis, these nonethylene receptor histidine kinases are stress-responsive and have a role in the regulation of plant response to abiotic stress such as drought and salt stress responses ^89^.

In plants, as in other eukaryotes, a diverse group of cell surface receptor-like protein kinases (RLKs) play a fundamental role in signal transduction processes. More than 600 genes belong to the receptor-like kinase (RLK) family. In Arabidopsis, among these RHS genes, RHS10, which encodes a proline-rich extensin-like receptor kinase (PERK), has a negative role in root hair elongation or tip growth ^90,91^. In the early steps of osmotic-stress signalling, several RLKs localized to the plasma membrane are involved in different plant species and recently have been suggested to be involved in the turgor pressure perception ^92^.

The transcription factors group DEHYDRATION-RESPONSIVE ELEMENT-BINDING PROTEIN 2 (DREB2) contribute to stress tolerance by initiating transcription through the cis-element dehydration-responsive element (DRE) in response to stress stimuli and its expression is induced by heat shock, dehydration and high salinity ^93,94^. Overexpression of DREB2 isolated from lotus, improves salt tolerance in transgenic A.thaliana ^95^. Additionally, a comprehensive analysis of rice DREB2-type genes that encode transcription factors reported that it is involved in the expression of abiotic stress-responsive genes. Moreover, the over-expression of DREB2 led to higher salt resistance than that of the wild-type plants, higher germination rates and better root growth in rice ^96^.

The chloroplast serine-threonine protein kinase (STN7) is required for the phosphorylation of the light-harvesting system of photosystem II and for the state transitions in Arabidopsis. The state transitions is a process that allows a balance between photosystem II and photosystem I through light excitation energy in the photosynthetic machinery and thereby optimizing the photosynthetic yield ^97^. Furthermore, it operates in retrograde signalling through controlling redox balance in the electron transfer chain and in short term responses via phosphorylation of a thylakoid bound phosphoprotein ^98,99^.

The secretory pathway is of a vital importance in most eukaryotic cells and has an essential role in a large variety of bioactive molecules synthesis, transport and secretion. These molecules participate in intercellular communication ^100^. The transmembrane emp24 domain-containing protein (TMED)/p24 gene family contribute to the vesicular trafficking of proteins, Golgi dynamics, as well as intracellular protein trafficking ^101^. Furthermore, for some p24 family members, additional roles in the post Golgi compartments of the secretory pathway have been recently proposed ^100^ and it was reported to be upregulated under environmental stresses such as heat shock ^102^.

In the past few years, some CRKs (cysteine-rich receptor-like kinases) were reported to play a critical role in biotic and abiotic responses in Arabidopsis, such as ABA signaling, disease resistance, plant growth, cell death and response by extracellular ROS production ^103–105^. CRKs are required in rice NH1 (NPR1)-mediated immunity ^104^ and in Arabidopsis which is protected against apoplastic oxidative stress ^106^ also it enhanced its pattern-triggered immunity by being overexpressed ^105^. Additionally, some genes belong to CRKs family act as a positive regulator of plant tolerance to salt stress ^107^.

### DArT assay

Compared to co-dominant SSRs and SNPs markers, DArT is a bi-allelic dominant marker, therefore; provides less genetic information ^108^. On the other hand, due its easy development it could provide a viable alternative for estimating the relation between various genotypes ^109110^. DArT technology was successfully applied in population genetics in order to detect useful loci associated with salinity tolerance in barley ^35,111,112^ and bread wheat ^17,113^.

In this research, we have used 3031 DArT markers. The maximum value of PIC was 0.375 and the minimum was 0.01 with an average of 0.13, which implies a locus of moderate informativeness ^64^. On the other hand, the maximum dissimilarity revealed with DArT markers was 0.2 and the minimum was 0, while the median was 0.012. DArT markers clustered chickpea genotypes into two clusters where the first cluster was divided into three subclusters and the second cluster was divided in four different clusters where G128434 was clustered in one cluster (FIGURE 2). Compared to the SSR phylogenetic tree, DArT markers were more comprehensive and showed more variation between different genotypes. MLM GWAS analysis ^114^was used to detect salinity associated DArT markers. Only three DArT markers (DART2393, DART769 and DART2009) showed a significant score. According to UniProtKB (www.uniprot.org), the two genes belong to catalytic activity also they are involved in cellular metabolic processes. DArT markers, DART2009 and DART769 were close to HIS3 and HIBCH which are located on Chr7 (TABLE 7 and FIGURE 5).

**Table 7:** The SNPs and DArTs physical position and marker-trait association significance p-value.

| Marker | Chr | Position | P-value |
| --- | --- | --- | --- |
| DART2393 | 2 | 2770214 | 0.000855266 |
| SNP2021 | 7 | 14771612 | 0.001071408 |
| SNP1268 | 2 | 3934500 | 0.001626215 |
| SNP1451 | 1 | 22106194 | 0.001700108 |
| DART769 | 5 | 19859119 | 0.003068851 |
| SNP1487 | 1 | 6383602 | 0.003249102 |
| SNP1667 | 3 | 39215537 | 0.003949766 |
| SNP2095 | 7 | 13746706 | 0.004381146 |
| SNP190 | 3 | 14518694 | 0.004515574 |
| SNP2247 | 2 | 33974127 | 0.005002969 |
| SNP1947 | 3 | 23327546 | 0.005224952 |
| DART2009 | 4 | 13717660 | 0.00557311 |
| SNP2331 | 4 | 45628179 | 0.006951469 |
| SNP948 | 4 | 16530791 | 0.008006447 |

Histidine is an essential dietary nutrient for animals, but it is synthesized de novo by plants and microorganisms. Imidazole glycerol-phosphate (IGP) dehydratase is involved in the histidine biosynthesis pathway. Thus, the pathway of histidine biosynthesis is a potential target for herbicide development ^115^.

3-hydroxyisobutyryl-CoA (HIBYL-CoA), a saline catabolite, has high activity toward isobutyryl-CoA ^116^ and its association with fatty acid biosynthesis or metabolism was also reported ^117^. The HIBYL-CoA is differentially expressed in the peanut roots in response to drought-responsive ^118^.

### SNP assay

SNP markers are more informative than classical molecular markers, it provides more comprehensive information about genetic mutations that is responsible for trait variability among different plant genotypes. SNP genotyping assay has been successfully used to dissect salinity tolerance in soybean ^25^ and cotton ^119^. In this research, 2500 SNPs markers were used for chickpea genotyping. The maximum PIC value was 0.0106 and the minimum was 0.58 with an average of 0.1634, which indicate a moderate loci informativeness. On the other hand, the maximum dissimilarity revealed by SNP markers was 0.53, the minimum was 0.004 and the median was 0.266. SNP-based phylogenetic tree clustered chickpea genotypes G70248 and G74929 in one cluster, while other genotypes were clustered in another major cluster. About 19 chickpea genotypes were extremely close to each other with moderately salinity tolerance and mostly from India and Pakistan (FIGURE 3).

Eleven SNP markers (TABLE 7) were associated with salinity tolerance in both filed and greenhouse. Ten SNP markers were close and have a “MODIFIER” effect to 16 chickpea genes (TABLE 6 and FIGURE 5). According to UniProtKB (www.uniprot.org), the gene ontology analysis showed that, at the molecular level these genes are involved in catalytic activity(3), binding(2) and trans-membrane transporter activity (1). While, on cellular component some shows a relationship to protein-containing complex (2),membrane part (3),intracellular organelle part, (2) and cell part (3). Additionally, on the biological process level some genes belong to metabolic process (1),cellular process (2), regulation of biological process (1) and localization (1). Some of these genes have a known direct or indirect impact on salinity tolerance in plants as follow;

NIPBL-like Plays an important role in the loading of the cohesin complex on to DNA and forms a heterodimeric complex (also known as cohesin loading complex) with MAU2/SCC4 which mediates the loading of the cohesin complex onto chromatin ^120,121^.

MAP-NDL is one of the major plant responses against salt stress that include microtubule depolymerization and reorganization, which is predicted to be pivotal for plant survival under abiotic stress ^122^. Microtubule organization is regulated by microtubule-associated proteins (MAPs) ^123^, of which MAP-NDL is a plant-specific protein that interacts with microtubules and regulates dynamics of microtubule ^122^ and may play a role in the anisotropic cell expansion and organ growth ^124,125^. Additionally MAP70-5, a divergent member of the MAP70 family of MAPs, is required for anisotropic cell growth in Arabidopsis and its overexpression causes epidermal cells to swell, induces right-handed organ twisting and maintain axial polarity to ensure the regular extension of plant organs ^124^.

P67 (a.k.a SUPPRESSOR OF VARIEGATION 7 (SVR7)) belongs to pentatricopeptide repeat (PPR) family, which contains 450 members and make up a significant proportion (6%) of the unknown functional proteins in Arabidopsis ^126^. They are defined by the presence of a canonical 35-amino-acid motif, repeated in tandem up to 30 times ^127^. The PPR proteins are considered to react with specific RNA in the cellular organelles and play a role in RNA processing or translation ^128^. It has been reported that they are involved in the splicing effects during the chloroplast development and the abiotic stress response in rice. A characterized mutant displays chlorotic striations in the early development, enhanced sensitivity to ABA and salinity and accumulation of more H2O2 than the wild-type ^129^. The Arabidopsis nuclear-localized PPR protein SVR7 is essential for the translation of the chloroplast ATP synthase subunits. SVR7 mutants were shown to accumulate higher levels of ROS and display increased sensitivity to H2O2 with decreased photosynthetic activity ^130^.

PUB9 belongs to ubiquitin (Ub) targeting proteins. It plays an important role in the degradation of proteins by the proteasome through polyubiquitination of substrate proteins via an enzyme cascade consisting of activating (E1), conjugating (E2) and ligating (E3) enzymes ^131,132^. It participates in several events in the life of the plant such as a hormone and biotic/abiotic stress signaling pathways ^133^. It functions as an E3 ubiquitin ligase and may be involved in the abscisic acid mediated signaling pathway at least during the germination stage ^134^. Its over-expression has been reported in root transcriptome analysis in grape genotypes with contrast translocation pattern of excess manganese from root to shoot ^135^.

Translocase of Outer Mitochondrial Membrane 6 (TOMM) is a component of the complex that is responsible for cytosolically synthesized mitochondrial pre-proteins ^136^. In Saccharomyces cerevisiae, seven TOMM genes associated with mitochondrial biosynthesis were significantly repressed after heat treatment ^137^.

Ubiquitination, the reversible protein conjugation with ubiquitin (Ub), is a post-translational modification that enables rapid and specific cellular responses to stimuli, without requirement of de novo protein synthesis. It has a central role in regulating many key cellular and physiological processes, including responses to biotic and abiotic stresses ^138,139^. The Ubiquitin-proteasome system (UPS) plays a central role in the efficient perception of different environmental stresses by suppressing stress signaling pathways during favorable growth conditions and thus, eliminating negative regulators of signaling responses to a stimulus ^140,141^.

Arabidopsis thaliana TRANSPARENT TESTA 12 (AtTT12) encodes a multidrug and toxic compound extrusion (MATE) transporter that contributes in seed coat pigmentation and functions as a vacuolar flavonoid/H+-antiporter that accumulates Proanthocyanidins in cells ^142,143^. Moreover, it has a role in the environmental stress tolerance ^144^. The upregulation of TESTA12 was reported in mechanisms of salt stress tolerance in R. stricta ^145^ and heat stress tolerance ^146^.

DNA-(apurinic or apyrimidinic site) lyase 2 (APEX2) works as a weak apurinic/apyrimidinic (AP) endodeoxyribonuclease in the DNA base excision repair (BER) pathway of DNA lesions that is induced by oxidative and alkylative agents. The presence of metal binding sites (MBS) is a typical feature of these proteins ^147,148^. The over-expression of AP has been thought to be responsible for enhancing osmotic stress tolerance in Medicago truncatula^148^. The AP lyase-dependent pathway to repair sites could generate new phenotypes and mutations ^149^. Additionally, it was reported that in Arabidopsis, the DNA glycosylase/lyase has an active role in DNA demethylation of ROS1, which possesses several enzymatic activities ^150^ and some AP genes function downstream of ROS1 in a ZDP-independent branch of the active DNA demethylation pathway in Arabidopsis ^151^.

MED4 is a mediator, which is a necessary component for RNA Pol II-mediated transcription of genes, where they play crucial roles in the basic process of transcriptional regulation of eukaryotic genes ^152^. In Arabidopsis, it was reported that some Med genes affect cell number and shoot meristem development ^153^, floral organ identity ^119^, embryo patterning and cotyledon organogenesis ^154^. Additionally, MEDs have been shown to play a role in biotic or abiotic responses, by the virtue of its ability to interact with several key transcriptional regulators ^155,156^. Moreover, in Arabidopsis, Med4 along with other Med genes showed more than 2-fold increase in their transcript abundance in response to the presence of NaCl in the media ^152^.

VPS39 (vam6/Vps39-like) is a complex, in which Vam6p/Vps39p stimulates guanine nucleotide exchange on small guanosine triphosphatase (GTPase) Vam4p/ Ypt7p and activates it, which in turn plays a vital role in tethering through the association with class CVps complex ^157^. it was upregulated and differentially expressed in secretome of TiK, the highly virulent T. indica 158.

Various key factors/regulators and transcription factors (TFs) play a critical role(s) towards regulating the gene expression patterns in response to stress conditions to cope with biotic and abiotic stresses ^159^. OsHBP1b TF belongs to the bZIP family and localized within the Saltol QTL, its expression is induced upon salinity treatment in seedlings, where it maintains chlorophyll content and improves the antioxidant machinery ^160^.

CHOLINE TRANSPORTER-LIKE1 (CTL1) is involved in choline transport process in addition, it functions in sieve plate and sieve pore formation in plants, where a mutation in this gene could cause several phenotypic abnormalities, including reduced pore density and altered pore structure in the sieve areas associated with impaired phloem function ^161^.

Kinesin-5 is a molecular motor protein that is essential in mitosis and exists in plant, animals and yeast ^162–164^. Plants possess a large repertoire of microtubule-based kinesin motor proteins. The distinctive inventory of plant kinesins suggests that kinesins have evolved to perform specialized functions in plants ^165^. Kinesin-5 is necessary for cortical microtubule organization and its loss severely compromises spindle structure and cytokinesis. Additionally, the kinesin-5 motor is crucial for mitosis in Arabidopsis roots and a mutant allele in this gene causes severe cell division defects in pollen development and embryogenesis ^166^.

### The relationship between SSR, DArT and SNP assays

Kinship analysis depending on DArT and SNP markers separated chickpea genotypes in two major clusters, where one cluster contain most of the chickpea genotypes (FIGURE 6). By collapsing the tree branches where they have the same similarity mean, they were collapsed in 16 branches (FIGURE 7)

**Figure 6:**
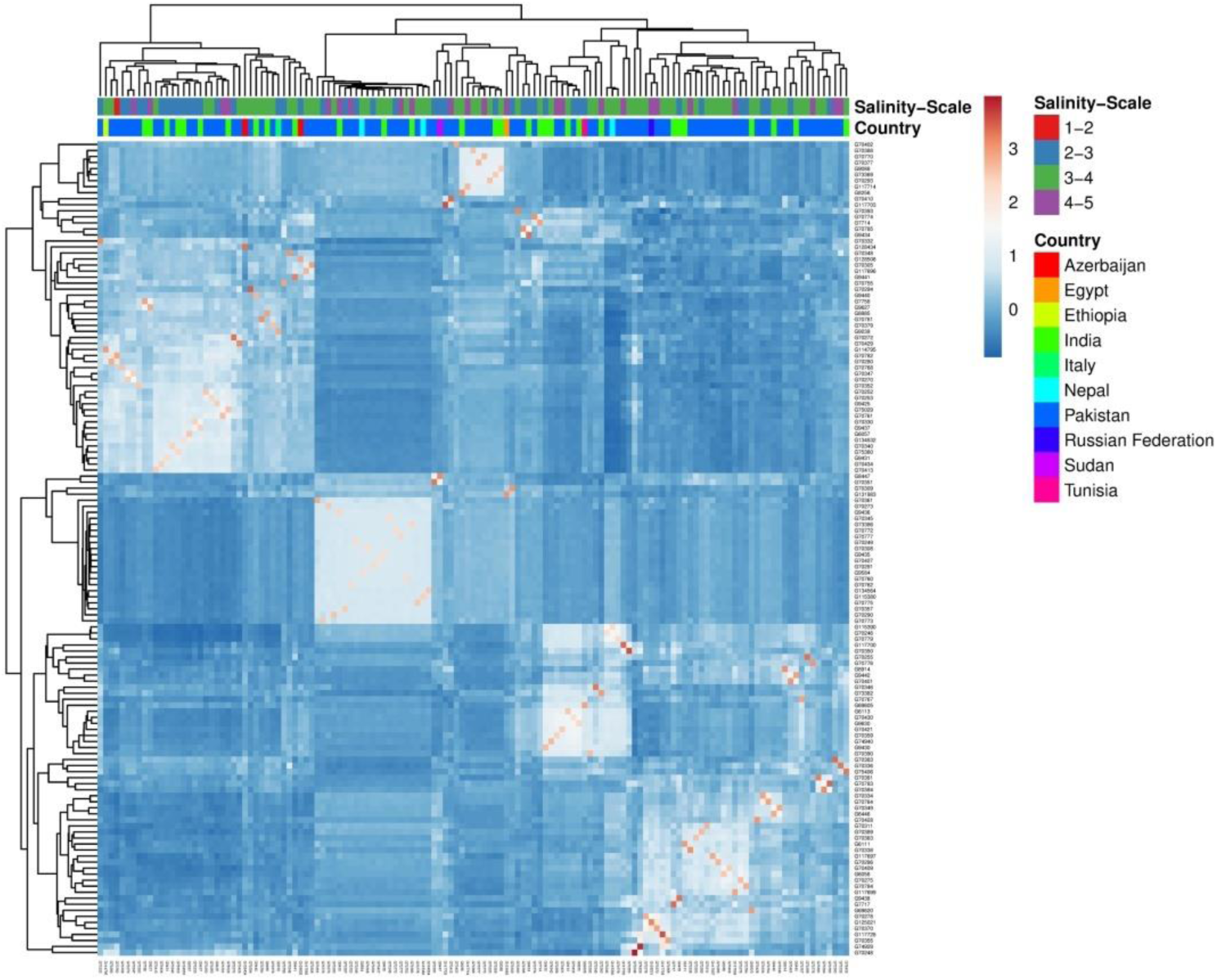
The kinship analysis for chickpea genotypes based on SSR, SNP and DArT markers.

**Figure 7:**
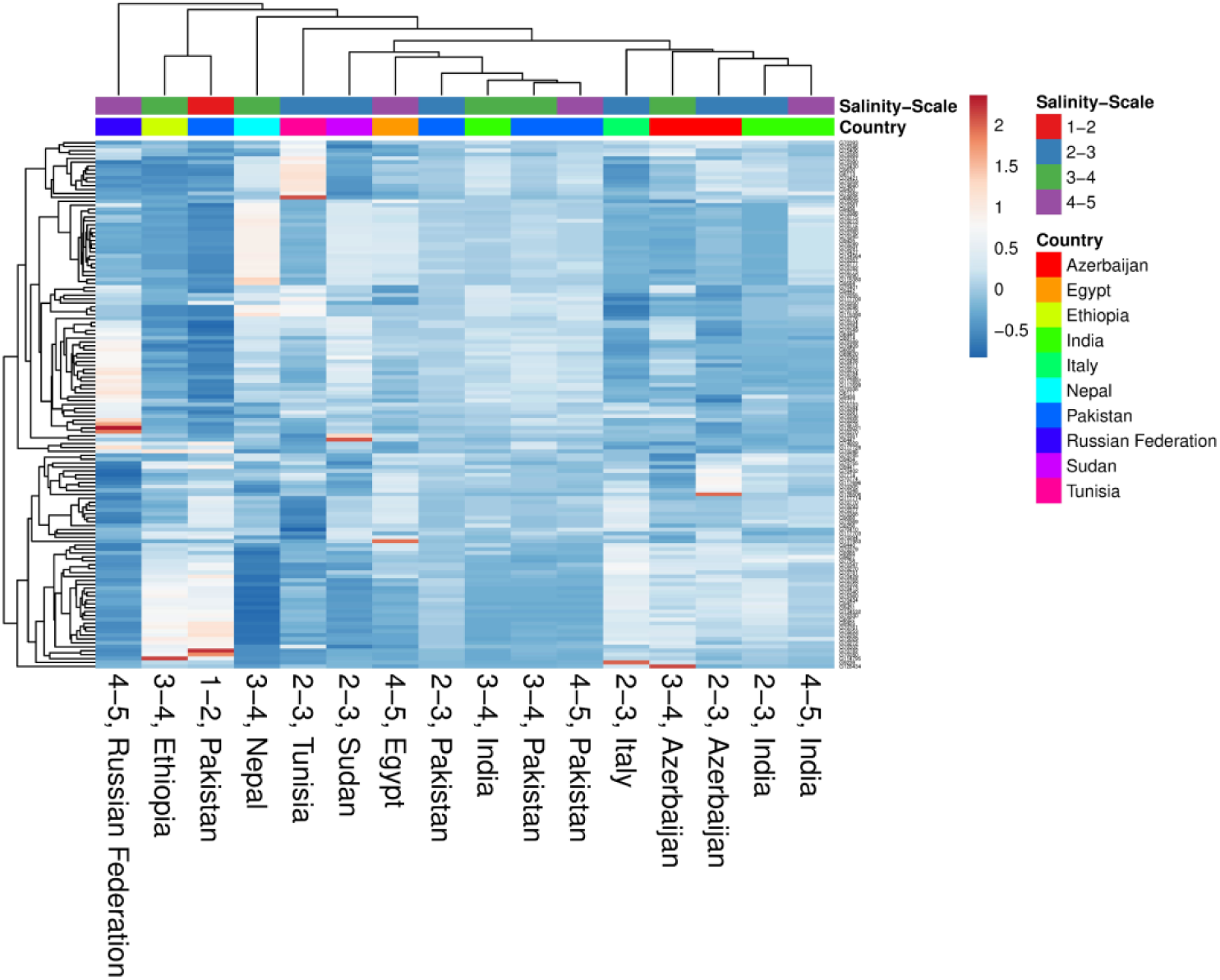
The kinship analysis for chickpea genotypes after collapsing branches with the same similarity mean based on SSR, SNP and DArT markers.

Mantel test as implemented in the GenAlex molecular analysis tool. It is employing 9999 random iterations which was used to calculate correlation matrices between the three marker systems that were used in this study. The correlation between SSR and both of DArT and SNP markers produced non or very low R^2^, which demonstrated a very weak correlation between the SSR markers and other markers. This could be due to the small number of SSR markers used in this study (FIGURE 8 and 9). However, a positive correlation between SNP and DArT assay was revealed by Mantel test with R^2^ of 0.58 (FIGURE 10). Similar results were reported by Baloch *et al*., ^167^in wheat, they spotted a correlation of 0.775 between DArT and SNP markers. Except for chromosomes 6 and 8, salinity tolerance associated genes were distributed across the chickpea genome (FIGURE 5).

**Figure 8:**
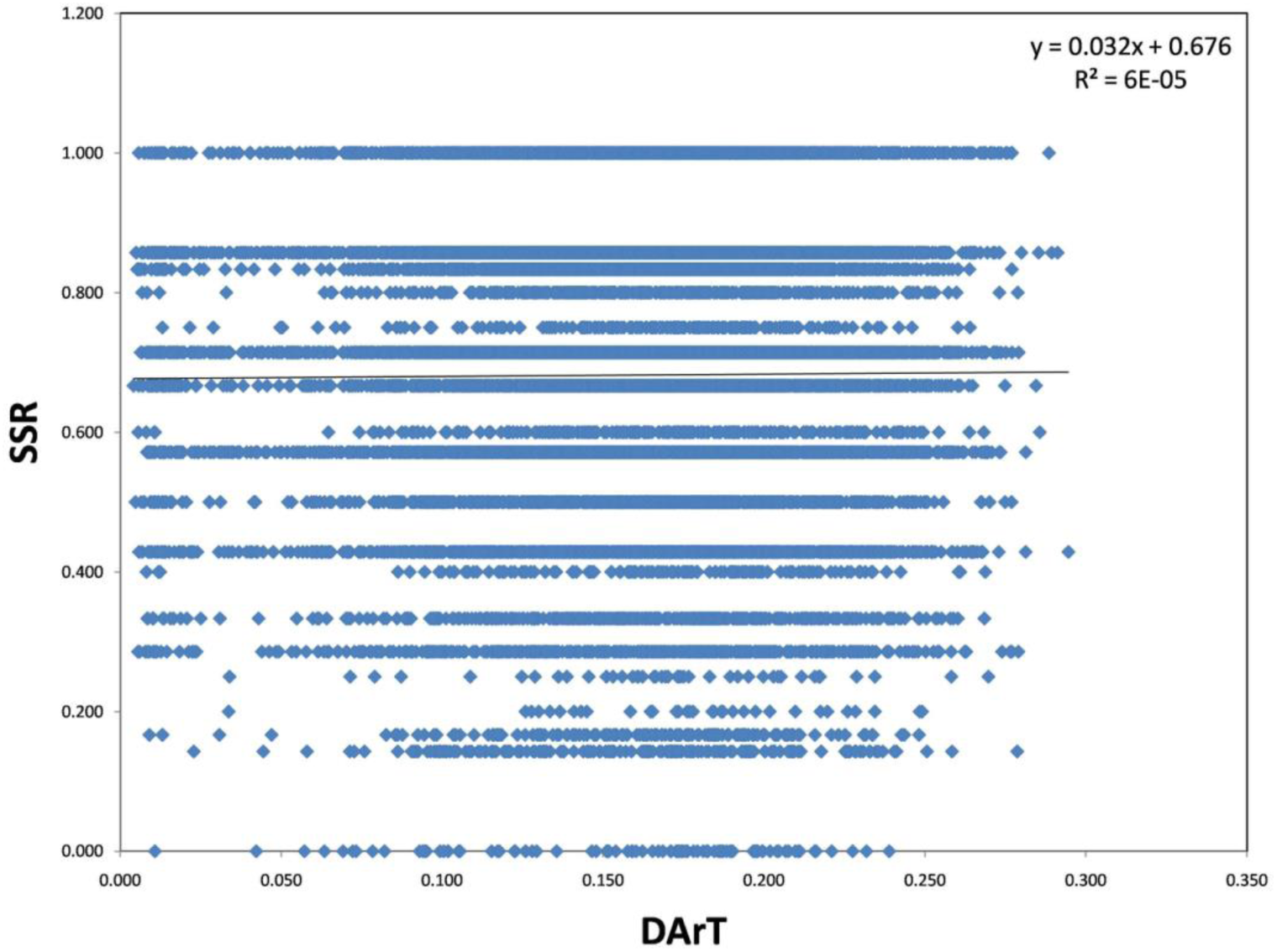
The mantel correlation between SSR and DArT markers.

**Figure 9:**
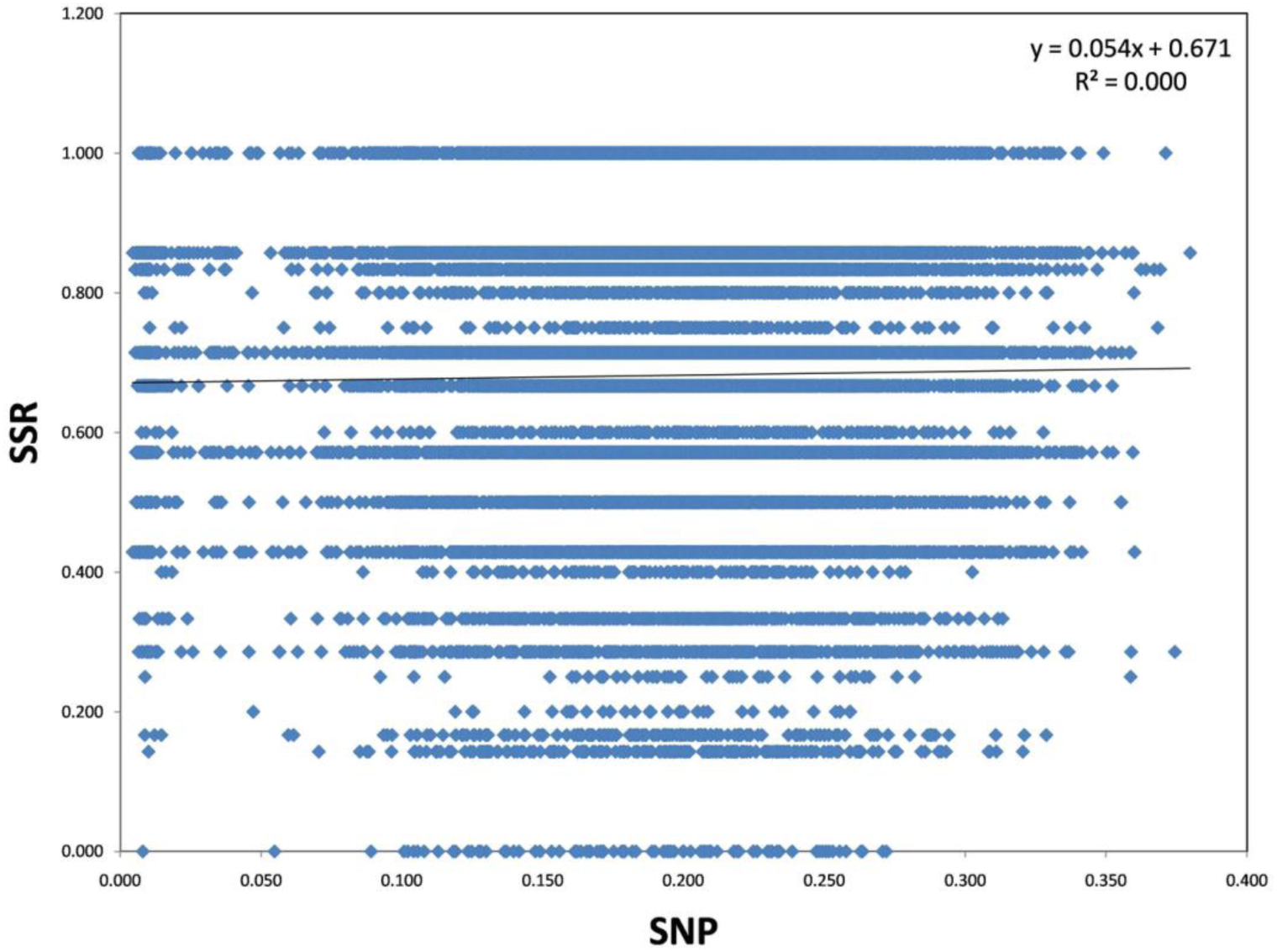
The mantel correlation between SSR and SNP markers.

**Figure 10:**
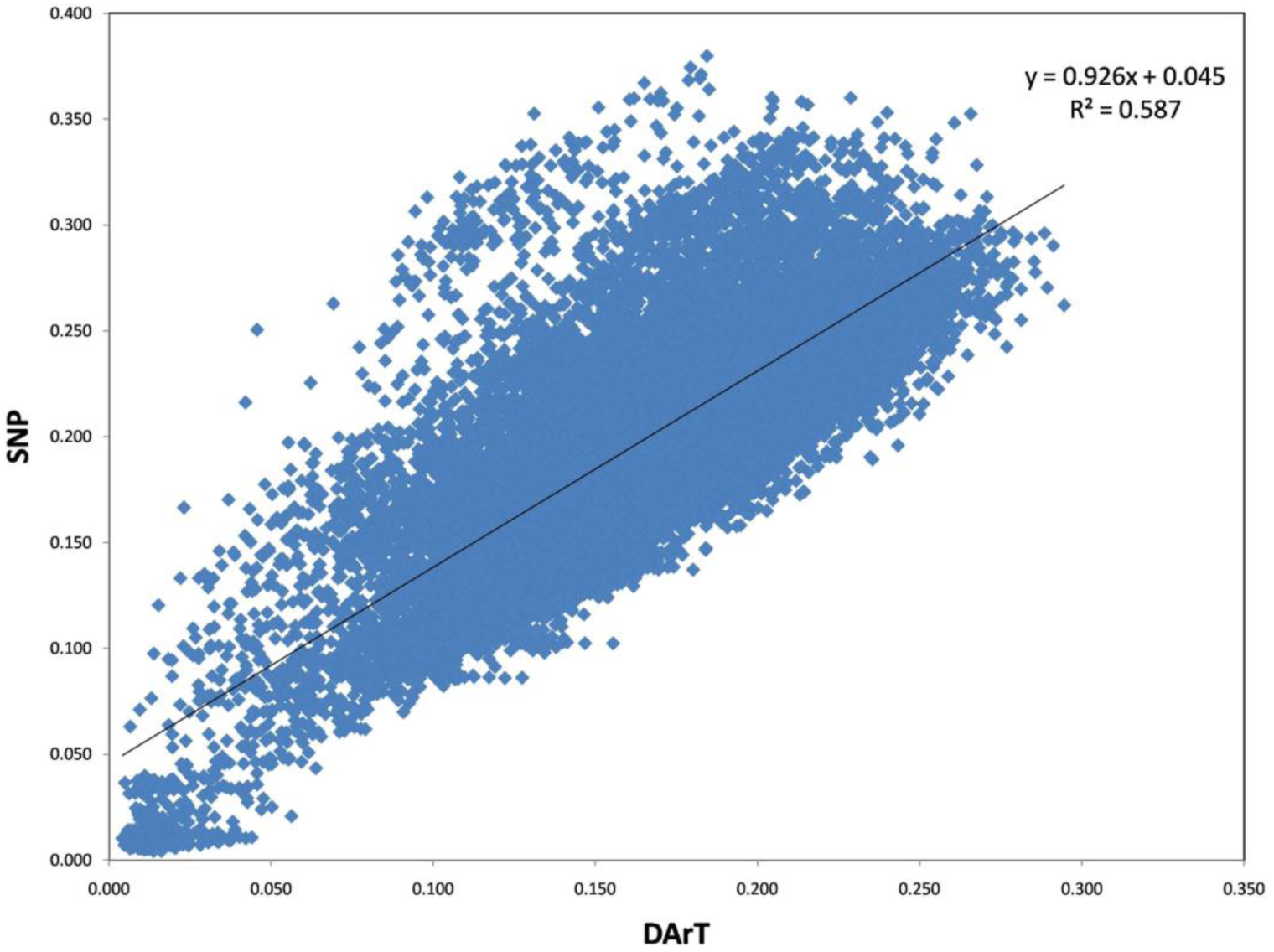
The mantel correlation between DArT and SNP markers.

## Acknowledgements

The authors gratefully acknowledge the financial support of International Center for Agricultural Research in the Dry Areas (ICARDA), Cairo, Egypt.

## Author Contributions

S.M.A, A.M.A and A.H. analyzed all experiments’ data and drafted the manuscript. S.M.A, M.H.M and H.A and conducted all greenhouse and filed experiments and the constitution of germplasm panel and its phenotyping. A.H, M.A.B, M.A.K and O.A.M participated in drafting and correcting the manuscript critically and gave the final approval of the version to be published. All authors have read and approved the final manuscript.

**Additional Information**

**Supplementary information** accompanies this paper at

### Competing Interests

The authors declare no competing interests.

